# Ribo-seq reveals IsrR-mediated translational repression of *SAOUHSC_02924* (*gabT*) during iron limitation in *Staphylococcus aureus*

**DOI:** 10.64898/2026.05.22.726194

**Authors:** Etornam Kofi Kumeko, Isabelle Hatin, Svetlana Chabelskaya, Enora Corler, Olivier Namy, Philippe Bouloc

**Affiliations:** Université Paris-Saclay, CEA, CNRS, Institute for Integrative Biology of the Cell (I2BC), 91198 Gif-sur-Yvette, France; Inserm, BRM [Bacterial RNAs and Medicine] - UMR_S1230, 35033 Rennes, France

## Abstract

Iron is essential for bacterial growth but can be toxic in excess. To maintain iron homeostasis, bacteria employ regulatory mechanisms, including small RNAs (sRNAs). In *Staphylococcus aureus*, we identified the sRNA IsrR as a critical mediator of the iron-sparing response, enabling bacterial fitness in iron-limited environments such as those encountered during host infection. Here, we use ribosome profiling (Ribo-seq) to define the translational regulatory network of IsrR under iron-limited conditions. Our analysis identifies multiple genes under IsrR control, including *SAOUHSC_02924* (*gabT*), which encodes a putative 4-aminobutyrate aminotransferase. Given that IsrR downregulates iron-dependent TCA cycle enzymes, we propose that repression of *gabT* prevents the accumulation of TCA cycle precursors under iron depletion, thereby avoiding metabolic imbalances. These findings expand the role of IsrR in metabolic reprogramming and highlight its contribution to *S. aureus* survival in iron-restricted host niches.

## INTRODUCTION

Iron is a necessary cofactor for enzymes involved in DNA synthesis, respiration, and other cellular functions. However, in excess, it contributes to oxidative stress by catalyzing the formation of reactive free radicals damaging cellular structures. Proper iron homeostasis is therefore essential to optimize fitness growth. In many bacteria, this regulation depends on the ferric uptake regulator (Fur), an iron-dependent transcriptional repressor, which controls the machinery required for iron acquisition (Steingard and Helmann, 2023).

As iron is required for pathogen growth, host sequesters iron as part of a defense mechanism known as nutritional immunity. Proteins such as transferrin and ferritin create an iron-depleted environment that limits the invasion of pathogens (Murdoch and Skaar, 2022). Central to the bacterial adaptation to iron scarcity is the iron-sparing response, a strategy that prioritizes cellular functions requiring less iron by reducing iron-intensive processes (Andrews et al., 2003). Bacterial small regulatory RNAs (sRNAs) are an integral part of this response, allowing bacteria to rapidly adjust gene expression at the post-transcriptional level. They typically act on specific messenger RNAs (mRNAs), often by binding near the ribosome-binding site (RBS), where they can inhibit translation by blocking ribosome access and possibly promote mRNA degradation. These mechanisms provide a rapid and flexible means of adjusting gene expression in response to environmental stressors, including during iron limitation (Wagner and Romby, 2015). In *Escherichia coli*, sRNA RyhB down-regulates the gene expression of iron-containing proteins in iron-restricted conditions to preserve the remaining iron for essential functions. (Masse and Arguin, 2005; Salvail and Masse, 2012). sRNAs with similar functions are present across distant bacterial species, underscoring the importance of iron regulation in bacterial growth (Chareyre and Mandin, 2018).

*Staphylococcus aureus*, an opportunistic human pathogen, can cause a wide range of infections, from mild skin infections to potentially fatal diseases such as endocarditis, sepsis, and pneumonia (Lowy, 1998). This adaptability to different host biotopes is in part due to its ability to counter host defenses, including nutritional immunity, enabling it to thrive even under metal ions-deprived conditions encountered in the host (Hammer and Skaar, 2011). We identified IsrR as the iron-<u>s</u>paring <u>r</u>esponse <u>r</u>egulator sRNA of *S. aureus*, demonstrating its importance for virulence and adaptation in iron-restricted growth conditions (Coronel-Tellez et al., 2022). IsrR base-pairs to Shine-Dalgarno (SD) sequences of several mRNAs encoding iron-sulfur-containing proteins, particularly with those associated with the dissimilatory nitrate reduction. Recently, two other enzymes containing iron-sulfur clusters were formally demonstrated to be downregulated by direct IsrR pairing to their corresponding mRNAs: MiaB, a tRNA-modifying enzyme (Barrault et al., 2024a) and CitB, the TCA cycle aconitase (Barrault et al., 2024b; Rios-Delgado et al., 2024); these results are also supported by global approaches investigating the *isrR* mutant (Ganske et al., 2024).

The present study investigates the regulatory role of IsrR employing ribosome profiling (Ribo-Seq). This powerful technique captures ribosome-protected RNA fragments providing a detailed snapshot of translational activity across the genome (Ingolia et al., 2009). By applying this technique to compare the isrR mutant and its isogenic parental strain under iron-replete and iron-restricted conditions, we evaluated the contribution of IsrR to the essential proteomic adjustments required for *S. aureus* survival in iron-limited environments. This analysis revealed the diversity of IsrR-mediated regulation and enabled the identification of novel IsrR targets. It provides a complementary insight into sRNA-dependent regulatory mechanisms enabling *S. aureus* to promote bacterial fitness under nutrient stress and consequently to withstand nutritional immunity.

## MATERIAL AND METHODS

### Bacterial strains, plasmids, and growth conditions

The bacterial strains and plasmids used in this study are listed in Supplementary Tables S1 and S2, respectively. All experiments were performed using the *S. aureus* HG003 strain (Herbert et al., 2010) and its derivatives (Supplementary Table S1). Gene annotations follow the NCTC8325 nomenclature (GenBank file CP00025.1) and were obtained from GenBank and Aureowiki (Fuchs et al., 2018).

Plasmids were constructed by Gibson assembly (Gibson et al., 2009) in *E. coli* IM08B (Monk et al., 2015) as outlined in Supplementary Table S2, using primers indicated in Supplementary Table S3 for PCR amplifications. Plasmid constructs were confirmed by DNA sequencing of the inserts and transferred into HG003 or its derivatives.

Chromosomal mutations—including point mutations, deletions, and insertions—were either obtained from prior studies or generated using *S. aureus*–*E. coli* shuttle vectors derived from pIMAY (Monk et al., 2012) or pRN112 (de Jong et al., 2017) as described (Table S1).

*E. coli* and *S. aureus* strains were grown in Lysogenic Broth (LB) and Brain Heart Infusion (BHI) media, respectively, as described (Barrault et al., 2024a). Antibiotics were used at the following concentrations: for *E. coli*, ampicillin (100 µg/mL) and chloramphenicol (20 µg/mL); for *S. aureus*, chloramphenicol (5 µg/mL), erythromycin (1 µg/mL), and kanamycin (60 µg/mL). Iron-limited conditions were achieved by supplementing media with 0.5 mM 2,2′-dipyridyl (DIP) 30 min prior to inoculation. This concentration moderately impacts *S. aureus* growth and reveals a reproducible difference between HG003 and its isogenic Δ*isrR* mutant (Barrault et al., 2024a).

### Ribosome profiling

Bacterial growth and sample preparation: *S. aureus* overnight cultures were cultivated in BHI broth at 37°C with shaking. Cultures were then diluted 1:1,000 into fresh BHI medium, supplemented with or without 2,2′-dipyridyl (DIP), and grown to an optical density at 600 nm (OD_600_) of 1.0. After rapid cooling on ice, cells were harvested by centrifugation (4,700 × *g*, 2 min, 4°C). Pellets were immediately flash-frozen in liquid nitrogen and stored at −70°C for downstream processing. Subsequent steps were performed as previously described (Chauhan et al., 2026).

### Biocomputing analysis

Pairing predictions between IsrR and the mRNA targets were made using intaRNA software (Mann et al., 2017), as described (Barrault et al., 2024a).

Analysis of ribosome profiling data: Ribosome profiling raw data were processed using the RiboDoc tool (v0.9.1.0) (Francois et al., 2021). The NCTC8325 reference genome (GenBank accession: CP00025.1) was used for alignment. Trimmed reads were filtered to retain ribosome-protected fragments (RPFs) ranging from 25 to 35 nucleotides in length. While read lengths of 28–30 nucleotides are typically optimal for P-site identification, the ideal length may vary depending on the sample and nuclease digestion method. Trimmed reads were filtered for ribosomal RNA (rRNA) by alignment against the 23S, 16S, and 5S rRNA sequences of *S. aureus* sourced from the SILVA database (Chuvochina et al., 2026) using Bowtie2 (Langmead and Salzberg, 2012); reads mapping to these references were excluded from downstream analyses. Reads were then mapped uniquely to mRNAs using both HISAT2 (Kim et al., 2019) and Bowtie2 (Langmead and Salzberg, 2012) with default parameters, allowing only one mismatch. The P-site offset was determined for each read length using the riboWaltz package (Lauria et al., 2018), which identifies the first base of each read based on signal alignment. To analyze the reading frame of ribosome-protected fragments (RPFs), each read was assigned a coordinate corresponding to the first nucleotide of the ribosome P-site, enabling the construction of metagene profiles reflecting the frequency of each footprint at every position along the transcript. Read counts were then quantified using HTSeq-count (Anders et al., 2015) in “intersection-strict” mode, restricted to coding sequence (CDS) regions to minimize noise from UTR regions and read pile-ups at initiation and termination sites. Differential gene expression analysis was subsequently performed with DESeq2 (Love et al., 2014) to assess variability between conditions. Genes were considered to show statistically significant expression changes when the adjusted p-value was below 0.01, with no threshold applied to the log2(fold change).

### Translational reporter assay for sRNA activity

Translational reporter fusions were constructed as described ((Barrault et al., 2024a); Tables S1–S2; Figure S1). In brief, the 5′ untranslated regions (5′UTRs) and initial codons of target mRNAs were fused in-frame to the coding sequence of the fluorescent protein mAmetrine. These fusions were placed under the control of the *sarA* P1 promoter (P_1_*sarA*). The constructs were assembled in two steps: first, the P_1_*sarA*-5′UTR-mAm fusions were generated using the pRN112 plasmid (de Jong et al., 2017), and then transferred into pIM-locus2 (Coronel-Tellez et al., 2022) for integration into the *S. aureus* chromosome of various strains (Table S1).

### Statistical tests

All data described in this paper originates from repeated experiments. Their number is indicated in the legend of figures (N = # of biological replicates). Error bars represent the standard deviation of N results. Statistical analyses between two groups were evaluated by a t-test using Excel. Relevant P-values were included in the figures using asterisk symbols (*, 0.05 ≤ p-value > 0.01; **, 0.01 ≤ p-xx).

## RESULTS

### Iron Depletion Triggers Translational Reprogramming in HG003

In Gram-negative bacteria, sRNAs primarily regulate gene expression by modulating mRNA stability, either promoting degradation or, less commonly, stabilization. As a result, transcriptomic comparisons are standard approaches for identifying sRNA targets. However, our previous transcriptome analysis of *S. aureus* HG003 and its Δ*isrR* mutant under iron-limited conditions revealed only minimal differences (Barrault et al., 2024a), suggesting that IsrR may not significantly alter mRNA abundance under these conditions. Similar results were observed in studies comparing transcriptomic and proteomic profiles following *isrR* deletion or overexpression (Ganske et al., 2024). Furthermore, the abundance and stability of known IsrR targets such as *fdhA, gltB2, miaB*, and *citB* mRNAs were largely unaffected by IsrR, despite confirmed base-pairing interactions (Barrault et al., 2024a; Barrault et al., 2024b; Coronel-Tellez et al., 2022). Together, these findings suggest that IsrR exerts regulatory control primarily at the level of translation, without significantly altering mRNA stability, at least under certain growth conditions.

To directly investigate whether IsrR regulates gene expression at the translational level, we employed genome-wide ribosome profiling (Ribo-seq), which quantifies ribosome occupancy on mRNAs. This approach allows us to capture active translation, independent of mRNA abundance. We first compared ribosome footprints in HG003 cultures grown in rich medium, either untreated or supplemented with the iron chelator 2,2′-dipyridyl (DIP). DIP treatment relieves Fur-mediated repression and induces endogenous *isrR* expression in the parental strain (Coronel-Tellez et al., 2022). For all Ribo-seq experiments, cultures were harvested at OD_600_ ≈ 1.0, and polysomes were isolated. Biological triplicates were performed for each condition, and changes were considered significant for fold change >2 and adjusted p < 0.01. A schematic overview of the Ribo-seq workflow and volcano plots summarizing the results is presented in Figure 1.

**Figure 1.**
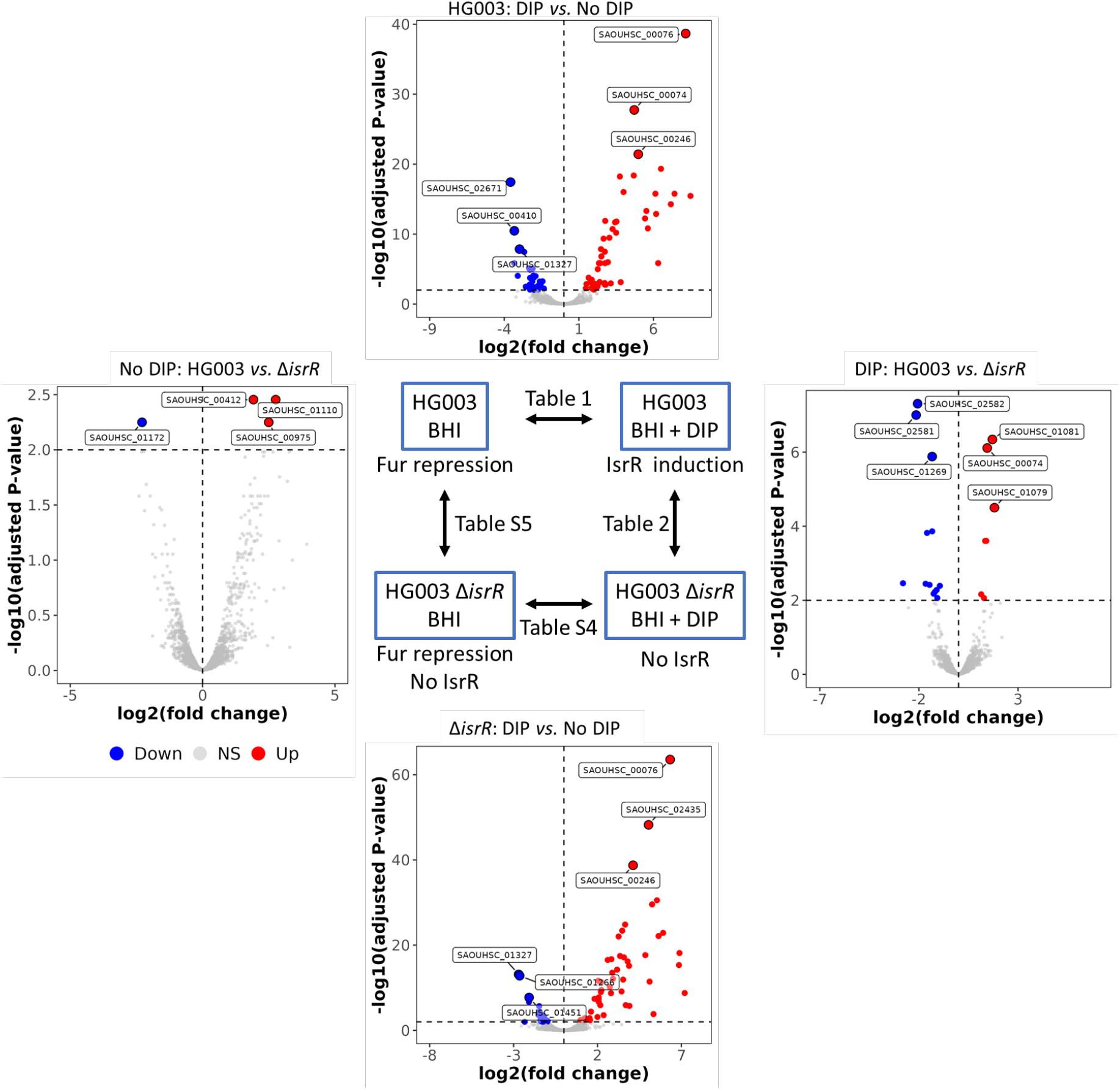
Schematic overview of Ribo-seq sample collection and volcano plots summarizing comparative results. Two bacterial strains, HG003 and its Δ*isrR* mutant derivative, were analyzed under two conditions: BHI medium without or with the iron chelator DIP. Cultures were sampled at an optical density (OD_600_) of 1.0. Comparative analyses are presented as follows: i) Effect of iron chelation (DIP) on the parental strain HG003 (Table 1); ii) Effect of iron chelation (DIP) on the Δ*isrR* mutant strain (Table S4); iii) Impact of IsrR deletion in iron-starved conditions in presence of DIP (Table 2) and iv) Impact of IsrR deletion in iron-replete conditions (Table S5). Volcano plots (x-axis: log_2_(fold change); y-axis: –log_10_(adjusted P-value)) illustrate the results for each table. Red dots: Significant read accumulation; Blue dots: Significant read reduction; Gray dots: No significant differences.

Ribo-seq analysis identified 51 genes with significantly increased translation in HG003 following DIP treatment (Table 1). Notably, 36 of these genes (71%) have been previously characterized as Fur-regulated (Fuchs et al., 2018; Horsburgh et al., 2001; Mazmanian et al., 2003), consistent with DIP’s role in relieving Fur repression. The remaining 15 genes included *cntA* and *cntB*, which encode components of the staphylopine transporter and are known to play a role in iron metabolism (Ghssein et al., 2016). Additionally, three loci (*SAOUHSC_00246, SAOUHSC_00247*, and *SAOUHSC_02873*) were upregulated; these have been previously reported as co-regulated with Fur targets (Casabona et al., 2017; Posada et al., 2014), suggesting a potential link to iron homeostasis. The operon comprising *SAOUHSC_00304, gcvH-L, sirTM*, and *lplA2* was also upregulated. This operon has been recently reported to be positively regulated by IsrR through a putative base-pairing interaction (Ganske et al., 2024).

**Table 1.** DIP-induced translational reprogramming in *S. aureus* HG003 mapped by ribosome profiling.

| ID | name | regulator | Pathway or function | log2 Fold Change* | Padj** |
| --- | --- | --- | --- | --- | --- |
| • Upregulated in DIP |  |  |  |  |  |
| SAOUHSC_00071 | *sirC* | Fur | Staphyloferrin B transporter | 3.81 | 0.00074 |
| SAOUHSC_00072 | *sirB* | Fur | Staphyloferrin B transporter | 2.74 | 0.00167 |
| SAOUHSC_00074 | *sirA* | Fur | Staphyloferrin B transporter | 4.71 | 1.76E-28 |
| SAOUHSC_00075 | *sbnA* | Fur | Staphyloferrin B biosynthesis | 8.48 | 3.49E-16 |
| SAOUHSC_00076 | *sbnB* | Fur | Staphyloferrin B biosynthesis | 8.16 | 2.11E-39 |
| SAOUHSC_00077 | *sbnC* | Fur | Staphyloferrin B biosynthesis | 7.16 | 5.34E-15 |
| SAOUHSC_00078 | *sbnD* | Fur | Staphyloferrin B transport | 6.32 | 1.43E-06 |
| SAOUHSC_00079 | *sbnE* | Fur | Staphyloferrin B biosynthesis | 6.13 | 1.68E-16 |
| SAOUHSC_00080 | *sbnF* | Fur | Staphyloferrin B biosynthesis | 6.51 | 4.65E-20 |
| SAOUHSC_00081 | *sbnG* | Fur | Staphyloferrin B biosynthesis | 5.62 | 1.53E-11 |
| SAOUHSC_00082 | *sbnH* | Fur | Staphyloferrin B biosynthesis | 6.18 | 1.31E-13 |
| SAOUHSC_00083 | *sbnI* | Fur | Staphyloferrin B biosynthesis | 5.43 | 5.81E-13 |
| SAOUHSC_00130 | *isdI* | Fur | Heme-degrading monooxygenase | 2.73 | 0.00098 |
| SAOUHSC_00131 |  |  | Iron acquisition and metabolism | 2.80 | 0.00172 |
| SAOUHSC_00246 |  |  | Putative efflux pump | 4.98 | 3.80E-22 |
| SAOUHSC_00247 |  |  | Putative choloylglycine hydrolase | 3.52 | 1.60E-12 |
| **SAOUHSC_00304** |  |  | Putative monooxygenase | 2.74 | 3.28E-08 |
| **SAOUHSC_00305** | ***gcvH-L*** |  | GcvH-like protein | 1.78 | 0.000591 |
| **SAOUHSC_00306** |  |  | Putative ADP-ribose binding module | 2.06 | 0.001277 |
| **SAOUHSC_00307** | ***sirTM*** |  | Deacetylase-like protein | 2.05 | 0.001721 |
| **SAOUHSC_00308** | ***lplA2*** |  | Lipoate salvage I | 1.65 | 0.000169 |
| SAOUHSC_00652 | *fhiC* | Fur | Staphyloferrin transport | 2.51 | 1.53E-07 |
| SAOUHSC_00653 | *fhuB* | Fur | Ferric hydroxamate uptake | 1.90 | 0.001225 |
| SAOUHSC_00654 | *fhuG* | Fur | Ferric hydroxamate uptake | 2.36 | 1.43E-06 |
| SAOUHSC_00746 | *sstA* | Fur | Siderophore transporter | 2.35 | 1.43E-06 |
| SAOUHSC_00747 | *sstB* | Fur | Siderophore transporter | 3.26 | 1.90E-11 |
| SAOUHSC_00748 | *sstC* | Fur | Siderophore transporter | 2.43 | 1.43E-06 |
| SAOUHSC_00749 | *sstD* | Fur | Siderophore transporter | 2.27 | 1.03E-05 |
| SAOUHSC_00833 | *ntrA* | Fur | Cofactor biosynthesis | 2.48 | 1.41E-08 |
| SAOUHSC_01031 | *cydA* |  | Cytochrome bd menaquinol oxidase | 1.51 | 0.001399 |
| SAOUHSC_01079 | *isdB* | Fur | Heme uptake | 7.41 | 1.68E-16 |
| SAOUHSC_01081 | *isdA* | Fur | Heme uptake | 3.50 | 6.37E-11 |
| SAOUHSC_01082 | *isdC* | Fur | Heme uptake | 3.43 | 2.03E-12 |
| SAOUHSC_01085 | *isdE* | Fur | Heme uptake | 3.14 | 0.00110 |
| SAOUHSC_01089 | *isdG* | Fur | Heme degradation VI | 2.38 | 0.00082 |
| SAOUHSC_01843 | *isdH* | Fur | Heme uptake | 5.52 | 4.88E-14 |
| SAOUHSC_02110 |  |  | Putative DnaQ family exonuclease | 2.02 | 0.008620 |
| SAOUHSC_02412 |  |  | Predicted transmembrane helices | 2.25 | 0.003098 |
| SAOUHSC_02427 | *htsC* | Fur | Staphyloferrin A transport | 2.76 | 1.43E-06 |
| SAOUHSC_02428 | *htsB* | Fur | Staphyloferrin A transport | 2.95 | 1.03E-06 |
| SAOUHSC_02430 | *htsA* | Fur | Staphyloferrin A transport | 2.77 | 1.30E-12 |
| SAOUHSC_02433 | *sfnaC* | Fur sigB | Staphyloferrin A biosynthesis | 3.75 | 5.75E-19 |
| SAOUHSC_02434 | *sfnaB* | Fur sigB | Staphyloferrin A biosynthesis | 4.00 | 9.59E-17 |
| SAOUHSC_02435 | *sfnaA* | Fur | Staphyloferrin A transport | 4.68 | 4.20E-19 |
| SAOUHSC_02436 | *sfnaD* | Fur | Staphyloferrin A biosynthesis | 1.87 | 0.000337 |
| SAOUHSC_02653 |  | Fur | Putative acetyltransferase | 1.96 | 0.007478 |
| SAOUHSC_02654 | *iruO* | Fur | Ferredoxin-NADP reductase | 2.66 | 4.47E-10 |
| SAOUHSC_02766 | *cntB* |  | Staphylopine-metal transport | 2.39 | 0.00069 |
| SAOUHSC_02767 | *cntA* |  | Staphylopine-metal transport | 1.86 | 0.005299 |
| SAOUHSC_02873 | *copA* |  | Cation transporter ATPase | 3.05 | 3.31E-10 |
| SAOUHSC_00364 | *ahpF* | PerR | Alkyl hydroperoxide reductase | 1.47 | 0.005574 |
| • Downregulated in DIP |  |  |  |  |  |
| SAOUHSC_00410 | *zagA* | Zur | Putative metal chaperon | -3.32 | 3.37E-11 |
| SAOUHSC_00706 | *fruR* | FruR CcpA | Transcriptional repressor | -2.09 | 0.003339 |
| SAOUHSC_00871 | *dltC* |  | D-alanyl-carrier protein | -2.12 | 0.000604 |
| SAOUHSC_00999 | *qoxD* |  | Quinol oxidase subunit IV | -1.55 | 0.001092 |
| SAOUHSC_01067 | *ctaM* | ctaA? | Putative heme chaperone | -2.02 | 0.000169 |
| **SAOUHSC_01103** | ***sdhC*** | CcpA | Succinate DH subunit | -2.66 | 3.53E-08 |
| **SAOUHSC_01104** | ***sdhA*** | CcpA | Succinate DH subunit | -2.09 | 8.17E-06 |
| **SAOUHSC_01105** | ***sdhB*** | CcpA | Succinate DH subunit | -2.34 | 7.34E-06 |
| **SAOUHSC_01327** | ***kata*** | CodY PerR | Catalase | -2.98 | 1.41E-08 |
| **SAOUHSC_01347** | ***citB*** | CcpE | Aconitase (FeS) | -2.05 | 8.44E-05 |
| SAOUHSC_01456 |  |  | Iron-uptake factor PiuB | -2.30 | 0.001703 |
| SAOUHSC_01466 | *recU* |  | Endonuclease | -2.01 | 0.009339 |
| SAOUHSC_01689 | *rpsT* |  | 30S ribosomal protein S20 | -2.56 | 0.003326 |
| **SAOUHSC_01776** | ***hemA*** |  | glutamyl-tRNA reductase | -1.62 | 0.000604 |
| **SAOUHSC_01960** | ***hemY*** | PerR | Protoporphyrinogen oxidase | -1.57 | 0.001993 |
| SAOUHSC_02340 | *atpC* |  | F0F1 ATP synthase subunit epsilon | -1.43 | 0.000564 |
| SAOUHSC_02343 | *atpG* |  | F0F1 ATP synthase subunit gamma | -1.34 | 0.006061 |
| SAOUHSC_02489 | *infA* |  | Translation initiation factor IF-1 | -2.27 | 0.008620 |
| SAOUHSC_02503 | *rpsQ* |  | 30S ribosomal protein S17 | -2.22 | 0.005014 |
| SAOUHSC_02504 | *rpmC* |  | 50S ribosomal protein L29 | -1.74 | 0.008620 |
| **SAOUHSC_02582** | ***fdhA*** | sigB | Formate dehydrogenase subunit $\alpha$ | -1.90 | 0.000100 |
| SAOUHSC_02671 | ***narK*** | Rex NreC | Nitrate/nitrite transporter | -3.58 | 3.64E-18 |
| SAOUHSC_02681 | ***narG*** | Rex NreC | Nitrate reductase subunit alpha | -3.33 | 1.43E-06 |
| SAOUHSC_02683 | ***nasE*** | Rex NreC | Nitrite reductase subunit | -1.78 | 0.002858 |
| SAOUHSC_02684 | ***nasD*** | Rex NreC | Nitrite reductase subunit | -2.27 | 0.000183 |
| SAOUHSC_02685 | ***nirR*** | Rex NreC | Sirohydrochlorin ferrochelatase | -3.09 | 9.18E-05 |
| SAOUHSC_02700 | ***mdeA*** | Zur | Putative MFS-type transporter | -2.27 | 1.21E-05 |

Name, regulator, and pathway/function were extracted from Aureowiki (Fuchs et al., 2018). \*Fold change > 2, \*\* Padj < 0.01. IDs/names in bold indicate previously reported putative or demonstrated IsrR-dependent gene regulation (gene from the same operon were associated)

**Table 2.** IsrR-dependent translational reprogramming in *S. aureus* HG003 upon DIP treatment mapped by ribosome profiling.

| ID | name | regulator | Pathway or function | log2 Fold Change* | Padj** |
| --- | --- | --- | --- | --- | --- |
| **Upregulated in presence of IsrR** |  |  |  |  |  |
| SAOUHSC_00074 | *sirA* | Fur | Staphyloferrin B transporter subunit | 1.44 | 0.000001 |
| SAOUHSC_00304 |  |  | Putative monooxygenase | 1.40 | 0.000249 |
| SAOUHSC_00545 | *sdrD* |  | Fibrinogen-binding protein | 1.34 | 0.000249 |
| SAOUHSC_01079 | *isdB* | Fur | Heme uptake | 1.81 | 0.000032 |
| SAOUHSC_01081 | *isdA* | Fur | Heme uptake | 1.71 | 0.000000 |
| SAOUHSC_01082 | *isdC* | Fur | Heme uptake | 1.28 | 0.008704 |
| SAOUHSC_02767 | *cntA* | Zur Fur | Staphylopin-metal transport | 1.14 | 0.006917 |
| **Downregulated in presence of IsrR** |  |  |  |  |  |
| SAOUHSC_00410 | *zagA* | Zur | Cobalamine synthesis | -1.46 | 0.003835 |
| SAOUHSC_01002 | *qoxA* | GraR | quinol oxidase AA3 subunit II | -0.94 | 0.004081 |
| SAOUHSC_01269 | *miaB* |  | tRNA base modification | -1.33 | 0.000001 |
| SAOUHSC_01270 | *ricA* |  | Regulatory Fe-S containing complex | -1.66 | 0.003585 |
| SAOUHSC_01347 | *citB* | CcpE | Aconitase | -1.59 | 0.000151 |
| SAOUHSC_01437 | *cvfC* |  |  | -1.26 | 0.006645 |
| SAOUHSC_01960 | *hemY* | PerR | Protoporphyrinogen oxidase | -1.11 | 0.005271 |
| SAOUHSC_02572 |  | SigB | Hypothetical protein | -2.80 | 0.003474 |
| SAOUHSC_02581 |  | SigA SigB | Formate hydrogenase | -2.14 | 0.000000 |
| SAOUHSC_02582 | *fdhA* | SigA SigB | Formate dehydrogenase subunit $\alpha$ | -2.07 | 0.000000 |
| SAOUHSC_02600 |  |  |  | -1.07 | 0.008677 |
| SAOUHSC_02681 | *narG* | Rex NreC | Nitrate reductase subunit alpha | -1.33 | 0.000137 |
| SAOUHSC_02924 | *gabT* | CodY | 4-aminobutyrate aminotransferase | -1.20 | 0.005963 |

Name, regulator and pathway/function were extracted from Aureowiki (Fuchs et al., 2018). \*Fold change > 2, \*\* Padj < 0.01.

In contrast, 27 genes exhibited significantly reduced ribosome occupancy in HG003 following DIP treatment while Fur acts mainly as a repressor (Table 1). This group included established iron-regulated genes, such as *sdhA, sdhB, katA*, and *citB* (Hempel et al., 2011). Strikingly, six of these genes (*citB, fdhA, katA, narG, nasD*, and *sdhC*) are confirmed or putative IsrR targets (Barrault et al., 2024a; Barrault et al., 2024b; Coronel-Tellez et al., 2022; Ganske et al., 2024; Rios-Delgado et al., 2024). This result is in agreement with a direct role for IsrR in their translational repression. Other genes located in operons that include established IsrR-regulated genes, such as *nasE* and *nirR* (within the *nasD* operon), were also downregulated, suggesting that IsrR binding to *nasD* ribosome binding sites (RBSs) may suppress translation of downstream genes through translational coupling. Similarly, *sdhA, hemA*, and *hemY* found here downregulated upon DIP-treatment were all recently proposed to be IsrR targets (Ganske et al., 2024; Rios-Delgado et al., 2025).

Collectively, these findings support a model in which IsrR accumulation during iron starvation represses the translation of specific mRNAs, contributing to an iron-sparing response. Additionally, IsrR may activate the translation of selected genes that lie outside the direct Fur regulon, suggesting a broader role in metabolic adaptation.

### Shared and Distinct Responses Between HG003 and Δ*isrR* Derivative to Iron Depletion

To dissect the specific role of IsrR in the iron starvation response, we compared DIP-induced translational changes in the Δ*isrR* mutant (Table S4) to those observed in the parental HG003 strain (Table 1). Of the 51 genes upregulated by DIP in HG003, 38 (76%) were also upregulated in Δ*isrR*. This overlap is expected, as these genes are predominantly Fur-regulated rather than by IsrR. Discrepancies were observed, mostly involving genes in operons potentially affected by DIP but excluded from significance due to adjusted p-value thresholds.

Strikingly, only 3 of the 27 genes repressed by DIP in HG003 were also downregulated in the Δ*isrR* strain. This marked discrepancy underscores the central role of IsrR in mediating translational repression during iron depletion.

### Putative Direct Targets of IsrR Identified by Comparative Ribo-Seq

To identify genes more directly regulated by IsrR at the translational level, we compared ribosome occupancy between HG003 and Δ*isrR* strains. In the absence of DIP, only minor differences were detected (Table S5), consistent with the low expression of *isrR* under iron-replete conditions.

Under iron-depleted conditions (DIP treatment), seven genes exhibited increased ribosome occupancy in HG003 relative to Δ*isrR*, while 13 genes were decreased (Table 2).

Five of the seven upregulated genes are established Fur targets. Their increased expression likely reflects a compensatory increase in iron demand due to the impaired iron-sparing response in the absence of IsrR. Notably, *SAOUHSC_00304* (Ganske et al., 2024) and *SAOUHSC_00545* (*sdrD*) mRNAs showed increased translation in wild-type HG003 compared to Δ*isrR*, suggesting possible positive regulation by IsrR, presumably independent of Fur.

From the 13 genes with decreased ribosome occupancy in HG003 compared to Δ*isrR*, five are IsrR validated targets, *fdhA, miaB, narG*, and *citB* (Barrault et al., 2024a; Barrault et al., 2024b; Coronel-Tellez et al., 2022; Rios-Delgado et al., 2024). Two additional genes, *SAOUHSC_01270* (*ricA*) and *SAOUHSC_02581*, were also repressed in HG003. Although these genes lack obvious IsrR base-pairing sites, their location downstream of *miaB* and *fdhA* suggests that their repression may occur via translational coupling (Figure S2).

Other putative IsrR-repressed genes include *SAOUHSC_00410* (*zagA*), *SAOUHSC_01002* (*qoxA*), *SAOUHSC_01437* (*cvfC*), *hemY, SAOUHSC_02572, SAOUHSC_02600*, and *SAOUHSC_02924*.

### Genes Upregulated in the absence of IsrR. Identification and Validation of New Direct IsrR Targets

To identify novel direct targets of IsrR, we focused on mRNAs differentially translated between Δ*isrR* and HG003 under DIP treatment (Table 2), excluding known IsrR targets and Fur-regulated genes. For the remaining candidates, we used IntaRNA to predict potential IsrR-mRNA interactions (Mann et al., 2017). Predictions were based on sequences spanning the transcription start site (TSS), as defined by Emote (Prados et al., 2016), to 30 nucleotides downstream of the start codon (10 amino acids). For SAOUHSC_02600, the region extended to 90 nucleotides (30 amino acids) to capture potential regulatory elements further downstream. For genes without annotated TSSs, predictions were based on published datasets (Mader et al., 2016). IntaRNA predicted significant base-pairing interactions for *SAOUHSC_00304, SAOUHSC_00410, SAOUHSC_00545, SAOUHSC_01002, SAOUHSC_01437, SAOUHSC_01960, SAOUHSC_02600* and *SAOUHSC_02924* mRNAs, with 6 of these interactions mapping to or near the Shine-Dalgarno (SD) sequence (Figures 2A and S3). To experimentally validate these predictions, we constructed translational fusions comprising the 5′ untranslated region (UTR) and the first 10 codons of each candidate gene, placed under the control of the constitutive PsarA1 promoter (Figures 2B and S1). The fusions were integrated into the chromosome of HG003, Δ*isrR*, Δ*fur*, and Δ*isrR* Δ*fur* strains using a temperature-sensitive pIM-locus2 plasmid, targeting the ‘locus 2’ intergenic region between *SAOUHSC_03030* and *SAOUHSC_03031* (Coronel-Tellez et al., 2022).

**Figure 2.**
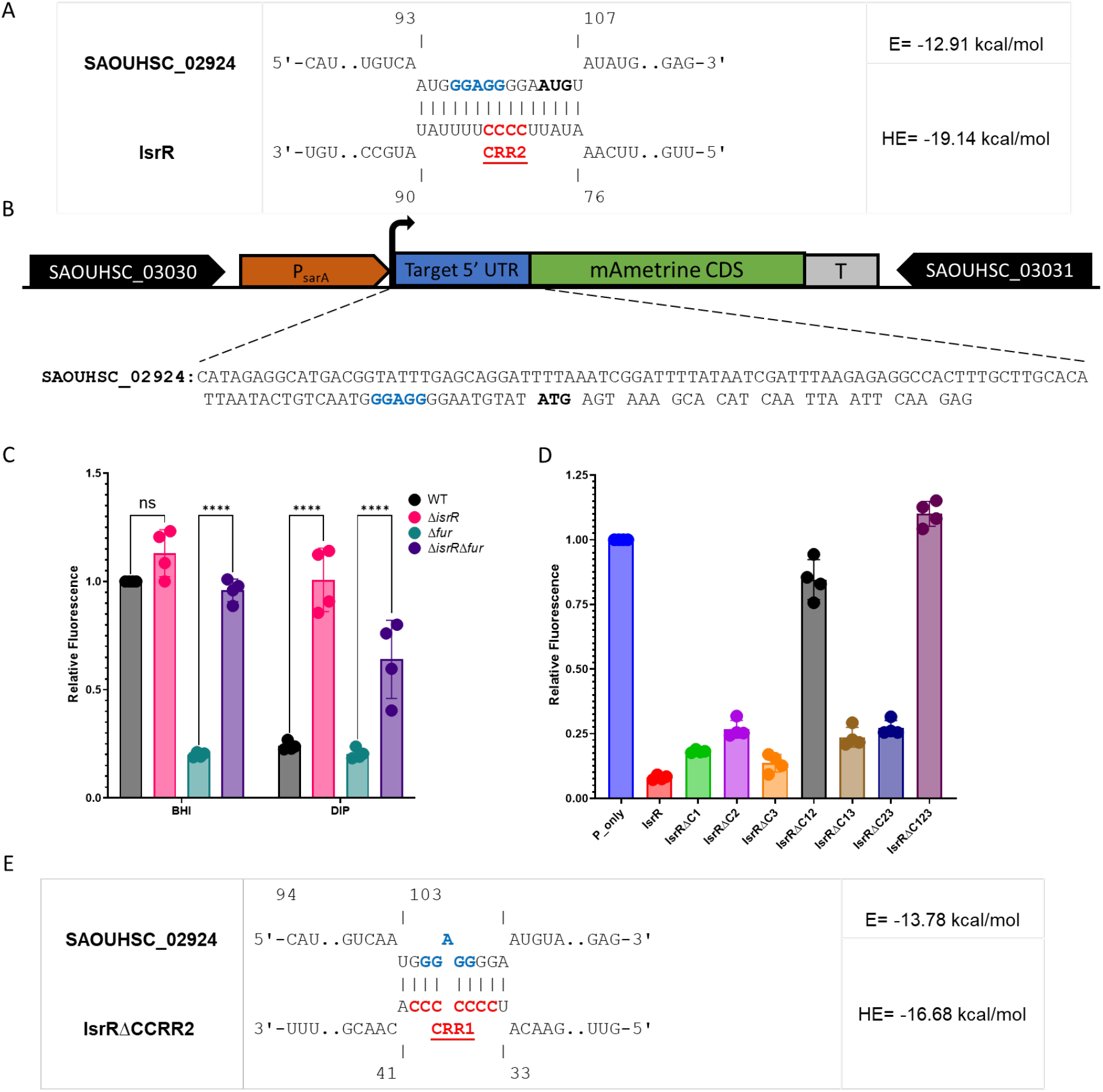
IsrR-mediated translational repression of the *SAOUHSC_02924* (*gabT*) reporter fusion. (A) Predicted interaction between the *SAOUHSC_02924* 5′ UTR (top strand) and the IsrR sRNA (bottom strand) as modeled by IntaRNA. The RBS is highlighted in blue, the ATG start codon in bold black, and the IsrR conserved regulatory region 1 (CRR1) in red. (B) Schematic of the chromosomal *SAOUHSC_02924*–mAmetrine post-transcriptional reporter fusion. The blue box (“Target 5′ UTR”) represents the 109-nucleotide 5′ UTR of *SAOUHSC_02924* and its first 10 codons. The corresponding sequence is shown below, with the RBS in blue and the ATG start codon in bold black. (C) IsrR-dependent repression of the *SAOUHSC_02924* reporter fusion. The *SAOUHSC_02924*–mAmetrine reporter (panel B) was integrated ectopically into the chromosome of *S. aureus* HG003 and its isogenic mutants (Δ*isrR*, Δ*fur*, and Δ*isrR* Δ*fur*). Cultures were grown in BHI medium with or without DIP supplementation. Relative fluorescence (normalized to the parental strain in BHI) is shown as bar graphs (mean ± SD, n = 4). Note that IsrR is fully expressed in the absence of Fur or in the presence of DIP (Coronel-Tellez et al., 2022). (D) Repression of the *SAOUHSC_02924* reporter fusion requires intact IsrR CRR1 or CRR2. Experiments were performed as in (C) (n = 4) using the Δ*isrR* strain harboring the *SAOUHSC_02924* reporter and plasmids expressing the indicated IsrR variants. For IsrR expression levels, see (Barrault et al., 2024a; Coronel-Tellez et al., 2022). (E) Predicted interaction between the *SAOUHSC_02924* 5′ UTR (top strand) and the IsrRΔC2 sRNA (bottom strand) as modeled by IntaRNA allowing 5 nucleotides for base-pairing in the seed motif. For color and other codes, see Figure 2A legend.

Strains were cultured in rich medium, with or without DIP, to induce *isrR* expression as previously observed in iron-depleted or Δ*fur* conditions (Coronel-Tellez et al., 2022). Of the 8 fusions tested, only the SAOUHSC_02924 UTR construct exhibited an IsrR-dependent change in fluorescence (Figures 2C and S4). The lack of response in the remaining fusions may indicate that critical regulatory regions were not captured in the constructs, or that these genes are indirectly affected by IsrR through downstream metabolic or regulatory pathways. Based on these results, *SAOUHSC_02924* was selected for further investigation.

### *SAOUHSC_02924* (*gabT*) mRNA is a bona fide IsrR target

sRNAs in Gram-positive bacteria often contain one or more C-rich motifs (CRRs) (Geissmann et al., 2009). These motifs typically base-pair with G-rich sequences in the Shine-Dalgarno (SD) region of target mRNAs, blocking ribosome binding, and inhibiting translation initiation. The contribution of individual CRRs to post-transcriptional regulation can vary depending on the target mRNA (Rochat et al., 2018). IsrR contains three single-stranded CRRs (Coronel-Tellez et al., 2022), each of which appears to regulate distinct subsets of targets (Barrault et al., 2024b).

To determine which CRRs mediate *SAOUHSC_02924* (*gabT*) repression, we analyzed the reporter fusion in strains expressing *isrR* variants lacking individual or combinations of CRRs. The reporter strain, which carries a chromosomal deletion of *isrR*, was complemented with plasmids constitutively expressing either wild type *isrR* or CRR deletion variants. *SAOUHSC_02924* (*gabT*) reporter expression was quantified by fluorescence measurements. An *isrR* variant lacking all three CRRs (*isrR*ΔC1ΔC2ΔC3) served as a negative control. Strains were cultured in rich medium, with or without DIP (to mimic iron depletion), and fluorescence was measured following overnight growth. Expression of wild-type IsrR led to the greatest reduction in *SAOUHSC_02924 (gabT)* reporter fluorescence. Deletion of any single CRR (*isrR*ΔC1, isrRΔC2, or isrRΔC3) did not abolish repression (Figure 2D), indicating functional redundancy among the CRRs for regulation.

We next tested IsrR variants lacking pairs of CRRs (IsrRΔC1ΔC2, IsrRΔC1ΔC3, and IsrRΔC2ΔC3). Repression of the *SAOUHSC_02924* (*gabT*) reporter was lost only in the IsrRΔC1ΔC2 variant, while repression remained intact in strains lacking CRR1/CRR3 or CRR2/CRR3 (Figure 2D). These results demonstrate that either CRR1 or CRR2 is sufficient to mediate *SAOUHSC_02924* (*gabT*) repression. Indeed, in the absence of CRR2, a second pairing involving CRR1 is proposed by IntaRNA (Figure 2E).

The *gabT* mRNA was previously identified as a putative direct target of RsaE based on computational predictions using RNAup (Geissmann et al., 2009). To experimentally validate this prediction, we introduced the translational fusion reporter (containing the predicted RsaE interaction region) into a Δ*rsaE* mutant strain. Fluorescence levels were then compared to those of its isogenic *rsaE*^+^ derivative. No significant difference was observed (Figure S5), indicating that *gabT* mRNA is likely not subject to post-transcriptional regulation by RsaE under the conditions tested.

Collectively, our data establish *SAOUHSC_02924* (*gabT*) mRNA is a likely direct IsrR target: its translation is specifically and reproducibly repressed by IsrR in a CRR-dependent manner, consistent with the canonical mechanism of sRNA-mediated translational inhibition.

## DISCUSSION

### Metabolic Plasticity and Virulence of S. aureus Under Iron Limitation

The ability of *S. aureus* to thrive under iron-limited conditions underscores its remarkable metabolic plasticity and virulence potential. In response to iron starvation, *S. aureus* deploys an adaptive strategy, encompassing not only the upregulation of canonical iron acquisition systems but also the reprogramming of central metabolic pathways via RNA-mediated regulation. To dissect the translational regulatory network of the iron-sparing small RNA IsrR, we employed Ribo-seq, a powerful tool for uncovering the targets of bacterial sRNAs by directly measuring translation at the genome-wide level in the presence or absence of a given sRNA (Wang et al., 2015). This approach is particularly valuable for identifying sRNA targets that are regulated at the translational level with moderate or no consequences on mRNA stabilities, as is the case for Gram-positive sRNA/target interactions. By comparing ribosome footprint densities between wild-type and IsrR mutant strains, we pinpointed mRNAs whose translation is directly repressed or activated by IsrR, even in the absence of changes in transcript levels. This strategy was previously applied to identify targets of RyhB, a functional homolog of IsrR in *E. coli*, confirming known targets and uncovering novel ones (Wang et al., 2015). Notably, Ribo-seq approaches for both IsrR and RyhB led to the identification of *citB* (called *acnA* in *E. coli*) and *narG* mRNAs, highlighting conserved regulatory mechanisms.

### Validation and Expansion of IsrR Targets

Our Ribo-seq experiments validated previously characterized IsrR targets (*narG, fdhA, miaB*, and *citB* mRNAs), confirming the robustness of our approach (Barrault et al., 2024a; Barrault et al., 2024b; Coronel-Tellez et al., 2022; Rios-Delgado et al., 2024). A recent multi-approach study proposed 21 “very likely IsrR targets” (Ganske et al., 2024). Our analysis identified 20 genes whose translation rates are modulated by IsrR, with five (*hemY, fdhA, miaB, citB*, and *SAOUHSC_00304*) independently supported by both studies. Discrepancies between the two global analyses likely arise from differences in strains (HG003 vs. HG001), experimental conditions (growth media: BHI vs. TSB; DIP stress: 500 µM for 30 min vs. 600 µM for 1 h), and the stringency of thresholds applied in each study, potentially excluding some targets.

Of the 20 genes whose translation rates are altered by IsrR, several are known IsrR substrates or have strong supporting evidence for regulation (e.g., operon structure or Fur-dependent regulation; see Results). The remaining eight putative IsrR targets were tested using reporter fusions designed based on IntaRNA predictions. Only *SAOUHSC_02924* (*gabT*) showed results consistent with direct IsrR-dependent regulation. Further experiments confirmed that IsrR binds to two sites within the SD sequence of *SAOUHSC_02924* (*gabT*) mRNA, and that two of its three conserved regulatory regions can independently target *SAOUHSC_02924* (*gabT*) mRNA. This pattern—specifically involving CRR1 and CRR2—was also observed in the IsrR-mediated downregulation of *citB* mRNA translation using the same combination of CRR deletions (Barrault et al., 2024a). Notably, IsrR substrates linked to nitrate respiration require either both CRR1 and CRR2 (*e*.*g*., *fdhA* and *gltB* mRNAs) or CRR2 and CRR3 (e.g., *narG* and *nasD* mRNAs) (Coronel-Tellez et al., 2022). In contrast, repression of *miaB* mRNA, which encodes a tRNA-modifying enzyme, depends solely on CRR3 (Barrault et al., 2024b). These examples illustrate the adaptability of IsrR in modulating gene expression through distinct combinations of CRRs, highlighting the complexity of bacterial sRNA regulatory networks.

### Functional Implications of IsrR-Mediated Repression of *gabT*

The downregulation of *SAOUHSC_02924* (*gabT*) by IsrR during iron deficiency prompts the question: What is the physiological rationale for this regulation? In *E. coli*, GabT catalyzes the conversion of γ-aminobutyric acid (GABA) and 2-oxoglutarate into succinate semialdehyde and L-glutamate (Metzer et al., 1979). This reaction initiates the GABA shunt, a metabolic bypass that connects GABA catabolism to central carbon metabolism by circumventing the 2-oxoglutarate dehydrogenase step of the TCA cycle. In higher organisms, GABA aminotransferase contains a [2Fe-2S] cluster (Storici et al., 2004). However, bacterial orthologs lack the cysteine residues required for cluster coordination and do not bind iron. Thus, the downregulation of *SAOUHSC_02924* (*gabT*) by IsrR is unlikely to serve as a direct iron-sparing mechanism, unlike the regulation of iron-dependent enzymes such as CitB, MiaB, and others.

In *S. aureus, SAOUHSC_02924* (*gabT*) is conserved and considered part of the core metabolic machinery, contributing to the bacterium’s adaptability to environmental changes, including those encountered during host infection (Wu et al., 2025; Xian et al., 2021). It is repressed by CodY, a global transcriptional regulator that acts in response to nutrient availability, particularly branched-chain amino acids and GTP coordinating carbon and nitrogen metabolism with environmental conditions (Majerczyk et al., 2010; Pohl et al., 2009). GABA acts as both a carbon/nitrogen source and a key intermediate in the GABA shunt, which replenishes the TCA cycle via succinate. Bacteria acquire GABA either by importing it via the gamma-aminobutyrate permease (GabP) or by synthesizing it, primarily from L-glutamate via glutamic acid decarboxylase (Gad). Alternative pathways, such as those using ornithine or arginine, also contribute to GABA biosynthesis. Intracellular GABA is sequentially converted to succinic semialdehyde by GABA aminotransferase (GabT) and then to succinate by succinate semialdehyde dehydrogenase (SSDH). GabP, GabT, and SSDH are not universally conserved across bacteria. Comparative genomics suggest that in *S. aureus, SAOUHSC_01326, SAOUHSC_02924*, and *SAOUHSC_02363* encode putative GabP, GabT, and SSDH homologs, respectively. Given the absence of a conserved *gad* gene in *S. aureus*, intracellular GABA could originate from external sources. *SAOUHSC_02924* (GabT) may thus play a role in succinate production, linking GABA metabolism to the TCA cycle.

The succinate semialdehyde produced by GabT enzymes is typically oxidized to succinate by succinate semialdehyde dehydrogenase (GabD). However, *S. aureus* lacks a canonical *gabD* ortholog, suggesting that one of its six annotated semialdehyde dehydrogenases (COG1012) (Galperin et al., 2021) may fulfill this role. If the shunt is functional, it can feed succinate into the TCA cycle. Under iron limitation, many TCA cycle enzymes are impaired; repression of *SAOUHSC_02924* (*gabT*) by IsrR is therefore expected to prevent the wasteful accumulation of succinate and avoid maladaptive flux through a compromised TCA cycle.

### Broader Physiological Roles of the GABA Shunt

Beyond *S. aureus*, the GABA shunt plays diverse physiological roles across bacteria, fungi, and plants, contributing not only to metabolic flexibility but also to stress adaptation, redox balance, and virulence. For example, in *Cronobacter sakazakii*, deletion of *gabT* increases intracellular GABA accumulation, enhancing stress resistance and biofilm formation (Wu et al., 2025). Many bacteria also use the GABA shunt to manage acid stress: the glutamate decarboxylase (GAD) system produces GABA under acidic conditions, consuming protons, while GabT and downstream enzymes help maintain metabolic and redox balance (Feehily and Karatzas, 2013; Wu et al., 2025). In *S. aureus*, these broader functions suggest that *gabT* is not merely a metabolic accessory but an integral component of stress response and adaptation. Regulation of *gabT*—such as its downregulation under iron starvation via IsrR— may enable *S. aureus* to balance metabolic demands, prevent the accumulation of potentially harmful intermediates, and fine-tune virulence in response to host environments.

### IsrR as a Master Regulator of Metabolic Reprogramming

Our results demonstrate that IsrR coordinates a broader metabolic reprogramming than previously recognized, conserving iron and sustaining energy production in iron-depleted environments. The translational shifts observed in this study highlight the bacterium’s capacity to prioritize iron-sparing processes to maintain cellular homeostasis. By integrating Ribo-seq data with functional assays, we have expanded the repertoire of IsrR targets and revealed the intricate regulatory logic governing its activity.

## SUPPLEMENTARY DATA

Supplementary Data. Tables S1 to S5, figures S1 to S4.

## ACKNOWLEDGEMENT

We thank Patricia Kerboriou for her technical help. We thank our colleagues Nara Figureoa-Bossi, Lionello Bossi, Alexandra Gruss and Tchak Palo for helpful discussions and warm support. We acknowledge the sequencing and bioinformatics expertise of the I2BC High-throughput sequencing facility, supported by *France Génomique* (funded by the French National Program “*Investissement d’Avenir*” ANR-10-INBS-09).

## FUNDING

The research was supported by the *Agence Nationale de la Recherche* [ANR-19-CE12-0006 (sRNA-RRARE)]. EK was the recipient of a scholarship from the ‘SDSV’ (*Structure et dynamique des systèmes vivants*) doctoral school (*Université Paris-Saclay*) and from the *Fondation pour la Recherche Médicale* (FRM).

## CONFLICT OF INTEREST

The authors declare no competing interests.

## Notes

### Competing Interest Statement

The authors have declared no competing interest.

