## Supplemental data for "Ribo-seq reveals IsrR-mediated translational repression of *SAOUHSC_02924* (*gabT*) during iron limitation in *Staphylococcus aureus*"

### Table S1. *Staphylococcus aureus* strains

| Strain | Genotype | Construction/ Reference |
|---|---|---|
| **HG003** | *rsbU* and *tcaR* repaired, MSSA, agr+ | (Herbert et al., 2010) |
| **MJH010** | as NCTC8325-4 Δ*fur*::*tetR* | (Horsburgh et al., 2001) |
| **SAPhB1231** | as HG003 Δ*isrR*::tag135 | (Coronel-Tellez et al., 2022) |
| **SAPhB1542** | as HG003 Δ*fur*::*tetR* | (Coronel-Tellez et al., 2022) |
| **SAPhB1443** | as HG003 Δ*isrR*::*tag135* Δ*fur*::*tetR* | SAPhB1231 + phage φ80 grown on MJH010 |
| **Strains with 5'UTRs downstream the promoter P1$_{sarA}$ driving *mAm* translation** | | |
| **SAPhB2614** | as HG003 P1$_{sarA}$-5'UTR-*SAOUHSC_01002*::*mAm*** | HG003 + pLoc2-5'_01002-mAm |
| **SAPhB2616** | as HG003 Δ*isrR*::tag135 P1$_{sarA}$-5'UTR-*SAOUHSC_01002*::*mAm*** | SAPhB1231 + pLoc2-5'_01002-mAm |
| **SAPhB2618** | as HG003 Δ*fur*::*tetR* P1$_{sarA}$-5'UTR-*SAOUHSC_01002*::*mAm*** | SAPhB1542 + pLoc2-5'_01002-mAm |
| **SAPhB2620** | as HG003 Δ*isrR*::tag135 Δ*fur*::*tetR* P1$_{sarA}$-5'UTR-*SAOUHSC_01002*::*mAm*** | SAPhB1443 + pLoc2-5'_01002-mAm |
| **SAPhB2622** | as HG003 P1$_{sarA}$-5'UTR-*hemY*::*mAm*** | HG003 + pLoc2-5'hemY-mAm |
| **SAPhB2624** | as HG003 Δ*isrR*::tag135 P1$_{sarA}$-5'UTR-*hemY*::*mAm*** | SAPhB1231 + pLoc2-5'hemY-mAm |
| **SAPhB2626** | as HG003 Δ*fur*::*tetR* P1$_{sarA}$-5'UTR-*hemY*::*mAm*** | SAPhB1542 + pLoc2-5'hemY-mAm |
| **SAPhB2628** | as HG003 Δ*isrR*::tag135 Δ*fur*::*tetR* P1$_{sarA}$-5'UTR-*hemY*::*mAm*** | SAPhB1443 + pLoc2-5'hemY-mAm |
| **SAPhB2630** | as HG003 P1$_{sarA}$-5'UTR-SAOUHSC_02924::*mAm*** | HG003 + pLoc2-5'_02924-mAm |
| **SAPhB2632** | as HG003 Δ*isrR*::tag135 P1$_{sarA}$_5'UTR-*SAOUHSC_02924*::*mAm*** | SAPhB1231 + pLoc2-5'_02924-mAm |
| **SAPhB2634** | as HG003 Δ*fur::tetR* P1$_{sarA}$-5'UTR-*SAOUHSC_02924*::*mAm*** | SAPhB1542 + pLoc2-5'_02924-mAm |
| **SAPhB2636** | as HG003 Δ*isrR*::tag135 Δ*fur*::*tetR* P1$_{sarA}$-5'UTR-*SAOUHSC_02924*::*mAm*** | SAPhB1443 + pLoc2-5'_02924-mAm |
| **SAPhB2679** | as HG003 P1$_{sarA}$-5'UTR-SAOUHSC_02600::*mAm*** | HG003 + pLoc2-5'_02600-mAm |
| **SAPhB2681** | as HG003 Δ*isrR*::tag135 P1$_{sarA}$_5'UTR-*SAOUHSC_02600*::*mAm*** | SAPhB1231 + pLoc2-5'_02600-mAm |
| **SAPhB2683** | as HG003 Δ*fur::tetR* P1$_{sarA}$-5'UTR-*SAOUHSC_02600*::*mAm*** | SAPhB1542 + pLoc2-5'_02600-mAm |
| **SAPhB2685** | as HG003 Δ*isrR*::tag135 Δ*fur*::*tetR* P1$_{sarA}$-5'UTR-*SAOUHSC_02600*::*mAm*** | SAPhB1443 + pLoc2-5'_02600-mAm |
| **SAPhB2694** | as HG003 Δ*rsaE*::*tag* P1$_{sarA}$-5'UTR-*SAOUHSC_02924*::*mAm*** | SAPhB2630 + pIMAY- Δ*rsaE* |
| **SAPhB2696** | as HG003 Δ*isrR*::tag135 Δ*rsaE*::*tag* P1$_{sarA}$-5'UTR-*SAOUHSC_02924*::*mAm*** | SAPhB2632 + pIMAY- Δ*rsaE* |
| **SAPhB2716** | as HG003 P1$_{sarA}$_5'UTR-*cvfC*::*mAm*** | HG003 + pLoc2-5'cvfC-mAm |

| | | |
|---|---|---|
| **SAPhB2718** | as HG003 Δ*isrR*::tag135 $P1_{sarA}$_5'UTR-*cvfC*::*mAm** | SAPhB1231 + pLoc2-5'cvfC-mAm |
| **SAPhB2720** | as HG003 Δ*fur::tetR* $P1_{sarA}$-5'UTR-*cvfC*::*mAm** | SAPhB1542 + pLoc2-5'cvfC-mAm |
| **SAPhB2722** | as HG003 Δ*isrR*::tag135 Δ*fur*::*tetR* $P1_{sarA}$-5'UTR-*cvfC*::*mAm** | SAPhB1443 + pLoc2-5'cvfC-mAm |
| **SAPhB2724** | as HG003 $P1_{sarA}$_5'UTR-*SAOUHSC_02600*::*mAm** | HG003 + pLoc2-5'_02600-mAm |
| **SAPhB2726** | as HG003 Δ*isrR*::tag135 $P1_{sarA}$_5'UTR-SAOUHSC_02600::*mAm** | SAPhB1231 + pLoc2-5'_02600-mAm |
| **SAPhB2728** | as HG003 Δ*fur*::*tetR* $P1_{sarA}$-5'UTR-*SAOUHSC_02600*::*mAm** | SAPhB1542 + pLoc2-5'_02600-mAm |
| **SAPhB2730** | as HG003 Δ*isrR*::tag135 Δ*fur*::*tetR* $P1_{sarA}$-5'UTR-*SAOUHSC_02600*::*mAm** | SAPhB1443 + pLoc2-5'_02600-mAm |
| **HG003 Δ*isrR* allele with the *gabT* reporter fusion complemented by different *isrR* alleles** | | |
| **SAPhB2658** | as HG003 Δ*isrR*::tag135 $P1_{sarA}$_5'UTR-*gabT*::*mAm** p_only | SAPhB2632 + p_only |
| **SAPhB2687** | as HG003 Δ*isrR*::tag135 $P1_{sarA}$_5'UTR-*gabT*::*mAm** pIsrR | SAPhB2632 + pIsrR |
| **SAPhB2644** | as HG003 Δ*isrR*::tag135 $P1_{sarA}$_5'UTR-*gabT*::*mAm** pIsrRΔC1 | SAPhB2632 + pIsrRΔC1 |
| **SAPhB2646** | as HG003 Δ*isrR*::tag135 $P1_{sarA}$_5'UTR-*gabT*::*mAm** pIsrRΔC2 | SAPhB2632 + pIsrRΔC2 |
| **SAPhB2648** | as HG003 Δ*isrR*::tag135 $P1_{sarA}$_5'UTR-*gabT*::*mAm** pIsrRΔC3 | SAPhB2632 + pIsrRΔC3 |
| **SAPhB2650** | as HG003 Δ*isrR*::tag135 $P1_{sarA}$_5'UTR-*gabT*::*mAm** pIsrRΔC1ΔC2 | SAPhB2632 + pIsrRΔC1ΔC2 |
| **SAPhB2652** | as HG003 Δ*isrR*::tag135 $P1_{sarA}$_5'UTR-*gabT*::*mAm** pIsrRΔC1ΔC3 | SAPhB2632 + pIsrRΔC1ΔC3 |
| **SAPhB2654** | as HG003 Δ*isrR*::tag135 $P1_{sarA}$_5'UTR-*gabT*::*mAm** pIsrRΔC2ΔC3 | SAPhB2632 + pIsrRΔC2ΔC3 |
| **SAPhB2656** | as HG003 Δ*isrR*::tag135 $P1_{sarA}$_5'UTR-*gabT*::*mAm** pIsrRΔC1ΔC3 | SAPhB2632 + pIsrRΔC1ΔC2ΔC3 |

* integrated between *SAOUHSC_03030* and *SAOUHSC_03031*

### Table S2. Plasmids

| Plasmids | Properties | Construction / reference |
|---|---|---|
| **pIMAY** | Shuttle rep(Ts) vector in *S. aureus* | (Monk et al., 2015) |
| **pIM-locus2** | pIMAY derivative plasmid for chromosomal DNA integration between the *SAOUHSC_03030* and *SAOUHSC_03031* genes. This intergenic region is defined as "locus 2". | (Coronel-Tellez et al., 2022) |
| **pRN112** | pJB28 derivative plasmid. Used here as an intermediate plasmid for the construction of reporter genes with the mAmetrique (*mAm*) gene encoding a fluorescent protein. | (de Jong et al., 2017) |
| **pRN112-5'_00304** | *PsarA-5'SAOUHSC_00304-mAm* gene fusion | 3388/3389 on HG003 + 2600/2926 on pRN112 |
| **pLoc2-5'_00304-mAm** | Chromosomal integration of the *PsarA-5'-SAOUHSC_00304-mAm* gene fusion between *SAOUHSC_03030* and *SAOUHSC_03031* | 2259/2260 on pIM-locus2 + 3541/3542 on pRN112 - 5'_00304 |
| **pRN112-5'sdrD** | *PsarA-5'sdrD-mAm* gene fusion | 3390/3391 on HG003 + 2600/2926 on pRN112 |
| **pLoc2-5'sdrD-mAm** | Chromosomal integration of the *PsarA-5'sdrD-mAm* gene fusion between *SAOUHSC_03030* and *SAOUHSC_03031* | 2259/2260 on pIM-locus2 + 3541/3542 on pRN112 - 5'sdrD |
| **pRN112-5'qoxA** | *PsarA*- 5'qoxA-*mAm* gene fusion | 3394/3395 on HG003 + 2600/2926 on pRN112 |
| **pLoc2-5'qoxA-mAm** | Chromosomal integration of the *PsarA*- 5'qoxA-*mAm* gene fusion between *SAOUHSC_03030* and *SAOUHSC_03031* | 2259/2260 on pIM-locus2 + 3541/3542 on pRN112 - 5'qoxA |
| **pRN112-5'hemY** | *PsarA*- 5'hemY *mAm* gene fusion | 3396/3397 on HG003 + 2600/2926 on pRN112 |
| **pLoc2-5'hemY-mAm** | Chromosomal integration of the *PsarA*- 5'hemY *mAm* gene fusion between *SAOUHSC_03030* and *SAOUHSC_03031* | 2259/2260 on pIM-locus2 + 3541/3542 on pRN112 - 5'hemY |
| **pRN112-5'_02600** | *PsarA-5'*SAOUHSC_02600-*mAm* gene fusion | 3402/3403 on HG003 + 2600/2926 on pRN112 |
| **pLoc2-5'_02600-mAm** | Chromosomal integration of the *PsarA-5'_02600-mAm* gene fusion between *SAOUHSC_03030* and *SAOUHSC_03031* | 2259/2260 on pIM-locus2 + 3541/3542 on pRN1125'_02600 |
| **pRN112-5'gabT** | *PsarA-5'gabT -mAm* gene fusion | 3398/3399 on HG003 + 2600/2926 on pRN112 |
| **pLoc2-5'gabT-mAm** | Chromosomal integration of the *PsarA- 5'gabT - mAm* gene fusion between *SAOUHSC_03030* and *SAOUHSC_03031* | 2259/2260 on pIM-locus2 + 3541/3542 on pRN112-5'gabT |
| **pIsrR** | pRMC2ΔR derivative. Constitutive expression of IsrR | (Coronel-Tellez et al., 2022) |
| **pIsrRΔC1** | pRMC2ΔR derivative. Constitutive expression of IsrRΔC1 | (Coronel-Tellez et al., 2022) |
| **pIsrRΔC2** | PRMC2ΔR derivative. Constitutive expression of IsrRΔC2 | (Coronel-Tellez et al., 2022) |
| **pIsrRΔC3** | pRMC2ΔR derivative. Constitutive expression of IsrRΔC3 | (Coronel-Tellez et al., 2022) |
| **pIsrRΔC1ΔC2** | pRMC2ΔR derivative. Constitutive expression of IsrRΔC1C2 | (Coronel-Tellez et al., 2022) |

| | | |
|---|---|---|
| **pIsrRΔC1ΔC3** | pRMC2ΔR derivative. Constitutive expression of IsrRΔC1C3 | (Barrault et al., 2024) |
| **pIsrRΔC2ΔC3** | pRMC2ΔR derivative. Constitutive expression of IsrRΔC2C3 | (Barrault et al., 2024) |
| **pIsrRC1ΔC2ΔC3** | pRMC2ΔR derivative. Constitutive expression of IsrRΔC1C2C3 | (Barrault et al., 2024) |

**Table S3. Primers**

| Name | Sequence |
|---|---|
| **2259** | TCACTGAAAATTTGTATAAAGATTTAAGTC |
| **2600** | TTAGTTAATTATAACTAATTAAAAATGAGAAGTAAAC |
| **2926** | CCATCGTTCAAATTTAGTTATG |
| **3388** | ttctcatttttaattagttataattaactaaGGGTATATGAAGAGGGAATGGTA |
| **3389** | aataattcttcaccttttgaaacGGCATAGTCTAATACGCTTAATTTAA |
| **3390** | ttctcatttttaattagttataattaactaaATACATTTTTGTATTTAAAACAATTG |
| **3391** | aataattcttcaccttttgaaacTATTGCCGTTTTATTTTCTCTG |
| **3394** | ttctcatttttaattagttataattaactaaCCCATAAGCCCTTGTAAATATTG |
| **3395** | aataattcttcaccttttgaaacTAATAGAAGCAAAGACTTAAATTTTG |
| **3396** | ttctcatttttaattagttataattaactaaGCCGAATACACATCCATTATTTATC |
| **3397** | aataattcttcaccttttgaaacCGCTCCTATAATAGCCACTG |
| **3398** | ttctcatttttaattagttataattaactaaCATAGAGGCATGACGGTATTTG |
| **3399** | aataattcttcaccttttgaaacCTCTTGAATTAATTGATGTGCTTTAC |
| **3402** | ttctcatttttaattagttataattaactaaAAGAGCGAAAGTGGGTGGAC |
| **3403** | aataattcttcaccttttgaaacCCAAGCTTCGTATACTAACTCTGG |
| **3541** | ATCCTTTAAATCATTGCGTGCTatcgcgattgcatgcctgcagg |
| **3542** | GACTTAAATCTTTATACAAATTTTCAGTGAatcgcgagctgcataaaaaacgccc |

**Table S4. DIP-induced translational reprogramming in *S. aureus* HG003 Δ*isrR* mapped by ribosome profiling.**

| ID | name | regulator | Pathway or function | log2 Fold Change* | Padj** |
|---|---|---|---|---|---|
| **Upregulated in DIP** | | | | | |
| ***SAOUHSC_00072*** | *sirB* | Fur | Staphyloferrin B transporter subunit | 5.33 | 0.000156 |
| ***SAOUHSC_00074*** | *sirA* | Fur | Staphyloferrin B transporter subunit | 3.25 | 9.50E-23 |
| ***SAOUHSC_00075*** | *sbnA* | Fur | Staphyloferrin B biosynthesis | 6.85 | 4.85E-16 |
| ***SAOUHSC_00076*** | *sbnB* | Fur | Staphyloferrin B biosynthesis | 6.32 | 2.75E-64 |
| ***SAOUHSC_00077*** | *sbnC* | Fur | Staphyloferrin B biosynthesis | 6.88 | 7.09E-19 |
| ***SAOUHSC_00078*** | *sbnD* | Fur | Staphyloferrin B transport | 3.90 | 1.84E-06 |
| ***SAOUHSC_00079*** | *sbnE* | Fur | Staphyloferrin B biosynthesis | 5.90 | 1.33E-23 |
| ***SAOUHSC_00080*** | *sbnF* | Fur | Staphyloferrin B biosynthesis | 5.25 | 2.73E-30 |
| ***SAOUHSC_00081*** | *sbnG* | Fur | Staphyloferrin B biosynthesis | 7.19 | 1.72E-09 |
| ***SAOUHSC_00082*** | *sbnH* | Fur | Staphyloferrin B biosynthesis | 5.63 | 7.34E-23 |
| ***SAOUHSC_00083*** | *sbnI* | Fur | Staphyloferrin B biosynthesis | 5.54 | 2.95E-31 |
| ***SAOUHSC_00130*** | *isdI* | Fur | Heme-degrading monooxygenase | 3.52 | 1.24E-12 |
| ***SAOUHSC_00131*** | | | Iron acquisition and metabolism | 3.68 | 1.23E-06 |
| ***SAOUHSC_00246*** | | | Putative efflux pump | 4.11 | 1.80E-39 |
| ***SAOUHSC_00247*** | | | Putative choloylglycine hydrolase | 3.64 | 1.48E-25 |
| *SAOUHSC_00364* | *aphF* | PerR | | 1.53 | 0.001378 |
| *SAOUHSC_00412* | *mpsA* | | NADH dehydrogenase subunit 5 | 2.07 | 1.93E-08 |
| *SAOUHSC_00413* | *mpsB* | | | 2.16 | 1.27E-06 |
| *SAOUHSC_00628* | *mnhD2* | sigB | Cation/$H^+$ antiporter subunit D | 1.19 | 0.006842 |
| *SAOUHSC_00632* | *mnhG2* | sigB | Cation/$H^+$ antiporter subunit G | 1.99 | 0.000739 |
| ***SAOUHSC_00652*** | *fhiC* | Fur | Staphyloferrin transport | 2.60 | 3.03E-17 |
| ***SAOUHSC_00653*** | *fhuB* | Fur | Ferric hydroxamate uptake | 1.82 | 4.03E-08 |
| ***SAOUHSC_00654*** | *fhuG* | Fur | Ferric hydroxamate uptake | 2.07 | 4.79E-08 |
| ***SAOUHSC_00746*** | *sstA* | Fur | Siderophore transporter | 3.16 | 5.71E-15 |
| ***SAOUHSC_00747*** | *sstB* | Fur | Siderophore transporter | 3.87 | 7.17E-16 |
| ***SAOUHSC_00748*** | *sstC* | Fur | Siderophore transporter | 3.79 | 6.46E-17 |
| ***SAOUHSC_00749*** | *sstD* | Fur | Siderophore transporter | 2.73 | 6.22E-11 |
| *SAOUHSC_00821* | | sigB | | 1.61 | 4.16E-05 |
| ***SAOUHSC_00833*** | *ntrA* | Fur | Cofactor biosynthesis | 2.22 | 4.68E-12 |
| *SAOUHSC_00848* | *sufD* | PerR | | 1.19 | 0.002270 |
| *SAOUHSC_00850* | *sufU* | PerR | | 1.24 | 0.001378 |
| *SAOUHSC_00851* | *sufB* | PerR | | 0.99 | 0.00338 |
| *SAOUHSC_01079* | *isdB* | Fur | Heme uptake | 5.10 | 3.65E-12 |
| *SAOUHSC_01081* | *isdA* | Fur | Heme uptake | 3.57 | 7.42E-18 |
| *SAOUHSC_01082* | *isdC* | Fur | Heme uptake | 2.92 | 7.93E-13 |
| ***SAOUHSC_01089*** | *isdG* | Fur | Heme degradation VI | 2.36 | 0.000274 |
| ***SAOUHSC_01843*** | *isdH* | Fur | Heme uptake | 4.85 | 2.19E-18 |
| *SAOUHSC_02246* | *fhuD1* | Fur | | -1.55 | 0.004165 |

| | | | | | |
|---|---|---|---|---|---|
| ***SAOUHSC_02427*** | *htsC* | Fur | Staphyloferrin A transport | 3.43 | 7.41E-10 |
| ***SAOUHSC_02428*** | *htsB* | Fur | Staphyloferrin A transport | 2.80 | 1.88E-09 |
| ***SAOUHSC_02430*** | *htsA* | Fur | Staphyloferrin A transport | -2.82 | 1.99E-17 |
| ***SAOUHSC_02433*** | *sfnaC* | Fur sigB | Staphyloferrin A biosynthesis | 3.34 | 3.51E-18 |
| ***SAOUHSC_02434*** | *sfnaB* | Fur sigB | Staphyloferrin A biosynthesis | 3.47 | 4.04E-24 |
| ***SAOUHSC_02435*** | *sfnaA* | Fur | Staphyloferrin A transport | 5.04 | 5.76E-49 |
| *SAOUHSC_02554* | *fhuD2* | Fur | Heme ABC transporter | 2.20 | 1.00E-09 |
| ***SAOUHSC_02653*** | | Fur | Putative acetyltransferase | 2.87 | 2.95E-14 |
| ***SAOUHSC_02654*** | *iruO* | Fur | Ferredoxin-NADP reductase | 2.05 | 2.29E-12 |
| ***SAOUHSC_02767*** | *cntA* | | Staphylopine-metal complex transport | 1.31 | 0.003297 |
| ***SAOUHSC_02873*** | *copA* | | Cation transporter E1-E2 family | 2.23 | 3.27E-10 |
| *SAOUHSC_02874* | *copZ* | | Cation transporter E1-E2 family | 2.07 | 2.10E-07 |
| ***Downregulated in DIP*** | | | | | |
| *SAOUHSC_00113* | *adhE* | Rex | Acetaldehyde-CoA/alcohol DH | -1.16 | 0.002079 |
| *SAOUHSC_00545* | *sdrD* | | Fibrinogen-binding protein SdrD | -1.38 | 0.000134 |
| *SAOUHSC_00608* | *adh1* | Rex | Alcohol dehydrogenase | -1.16 | 0.001184 |
| *SAOUHSC_00721* | *queC* | PreQ1 ribos. | 7-cyano-7-deazaguanine synthase | -1.52 | 0.000237 |
| *SAOUHSC_00808* | | | | -1.39 | 0.002779 |
| *SAOUHSC_00838* | | | | -1.08 | 0.009723 |
| *SAOUHSC_01266* | | | | -2.63 | 1.54E-13 |
| *SAOUHSC_01268* | | | | -1.36 | 0.000364 |
| *SAOUHSC_01287* | *glnA* | CodY GlnR | Glutamine synthetase | -1.21 | 0.001752 |
| ***SAOUHSC_01327*** | *katA* | CodY PerR | Catalase | -2.69 | 7.27E-14 |
| ***SAOUHSC_01347*** | *citB* | Fur CcpA | Aconitase (FeS) | -1.24 | 0.000274 |
| *SAOUHSC_01424* | *murG* | | Lipid I N-acetylglucosaminyltransferase | -1.22 | 0.007152 |
| *SAOUHSC_01440* | | PreQ1 ribos. | Putative preQ0 transporter | -2.33 | 0.009838 |
| *SAOUHSC_01451* | *ilvA1* | CodY | Threonine dehydratase | -2.07 | 1.93E-08 |
| *SAOUHSC_01653* | *sodA* | CodY | Superoxide dismutase | -1.46 | 4.54E-05 |
| ***SAOUHSC_01776*** | *hemA* | | Glutamyl-tRNA reductase | -1.50 | 0.000237 |
| *SAOUHSC_02012* | *sgtB* | sigB | Glycosyltransferase | -1.19 | 0.006143 |
| *SAOUHSC_02108* | *ftnA* | Fur PerR | Ferritin | -2.08 | 2.20E-07 |
| *SAOUHSC_02412* | | | | -1.38 | 0.006842 |
| *SAOUHSC_02542* | *moeA* | | Molybdopterin biosynthesis protein | -0.99 | 0.009214 |
| *SAOUHSC_02666* | | | Putative lipoprotein | -1.47 | 2.12E-06 |
| *SAOUHSC_02744* | *opuCA* | CodY | ABC transp. ATP-binding protein | -1.25 | 0.009723 |
| *SAOUHSC_02830* | *ddh* | Rex | D-lactate dehydrogenase | -0.96 | 0.005683 |
| *SAOUHSC_02947* | *cysJ* | CymR | Sulfite reductase protein subunit α | -1.09 | 0.003678 |
| *SAOUHSC_02972* | *isaB* | | Immunodominant antigen B | -1.12 | 0.003535 |

Name, regulators, and pathway/function were extracted from Aureowiki (Fuchs et al., 2018). *Fold change > 2, ** Padj < 0.01.

**Table S5. Translational differences between *S. aureus* HG003 and its Δ*isrR* isogenic derivative grown in rich medium mapped by ribosome profiling.**

| ID | name | regulator | Pathway or function | log2 Fold Change* | Padj** |
|---|---|---|---|---|---|
| **Upregulated in HG003** | | | | | |
| *SAOUHSC_00412* | *mpsA* | | NADH dehydrogenase subunit 5 | 1.92 | 0.003509 |
| *SAOUHSC_01110* | *ecb* | SaeR / RNAIII | fibrinogen-binding protein-like protein | 2.76 | 0.003509 |
| SAOUHSC_00975 | | | | 2.49 | 0.005631 |
| **Downregulated in HG003** | | | | | |
| *SAOUHSC_01172* | *pyrE* | pyrR leader | orotate phosphoribosyltransferase | -2.28 | 0.005631 |

Name, regulators and pathway/function were extracted from Aureowiki (Fuchs et al., 2018). *Fold change > 2; ** Padj < 0.01.

**Figure S1. Translational Reporter Assay for sRNA Activity**

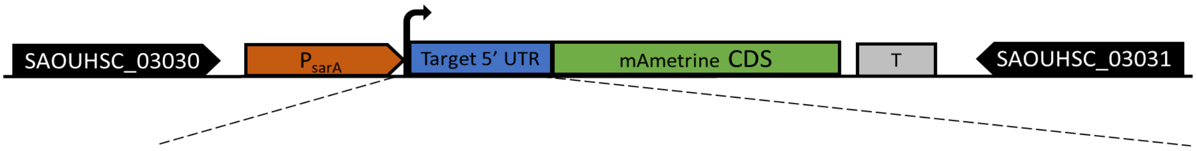

*SAOUHSC_00304*: GGGTATATGAAGAGGGAATGGTATTAAGAACGCTAAAATGTTATGTCGGTTTGACATGACAGGATAAGTTT**GGAGA**TGACGGA
**TTG** GTT AAA TTA AGC GTA TTA GAC TAT GCC
*SAOUHSC_00410*: AGAATTATTCCTATTTACAAATAAATTGAAGT**GAGGG**TATATAT ATG GCT AAA ATT CCA GTT ACG GTA TTA AGT
*SAOUHSC_00545*: ATACATTTTTGTATTTAAAACAATTGTATTATTTAAAAATTTGATGAATTA**GGAGTA**ATCTA ATG CTA AAC AGA GAA
AAT AAA ACG GCA ATA
*SAOUHSC_01002*: CCCATAAGCCCTTGTAAATATTGTGTAATAACTTAAAATTAATGTTAAGCCCTACATTTGTAGTATTA**GGAGG**TCAAAAAA GTG
TCA AAA TTT AAG TCT TTG CTT CTA TTA
*hemY*: GCCGAATACACATCCATTATTTATCGGTGCAATTATTGATGAAATCAAGTCTATATTTTAATGACGAAAGAAGCGTGAAACGTT
**GTG** ACT AAA TCA GTG GCT ATT ATA GGA GCG
SAOUHSC_02600: AAGAGCGAAAGTG**GGTGG**ACAACA GTG GCA AAA TTT AAT GTA GAG AAT GAA CAT

The schematic illustrates a chromosomal 5′UTR–mAmetrine post-transcriptional reporter fusion. The blue box represents the tested interaction region, comprising the 5′ untranslated region (UTR) of the target gene and its first 10 codons, fused in-frame with the coding sequence (CDS) of the fluorescent protein mAmetrine (green). This fusion is driven by the constitutive *PsarA* promoter (orange) and followed by a transcriptional terminator (grey).

The reporter fusions were integrated into the *S. aureus* chromosome at locus 2, between the genes *SAOUHSC_03030* and *SAOUHSC_03031*, using the integrative plasmid pIM-locus_2 (Coronel-Tellez et al., 2022). The tested sequences, including the ribosome-binding site (RBS, blue) and start codon (bold black), are detailed in the lower panel. *SAOUHSC_02924* is presented in Figure 1A.

**Figure S2. Operonic organization supporting translational coupling**

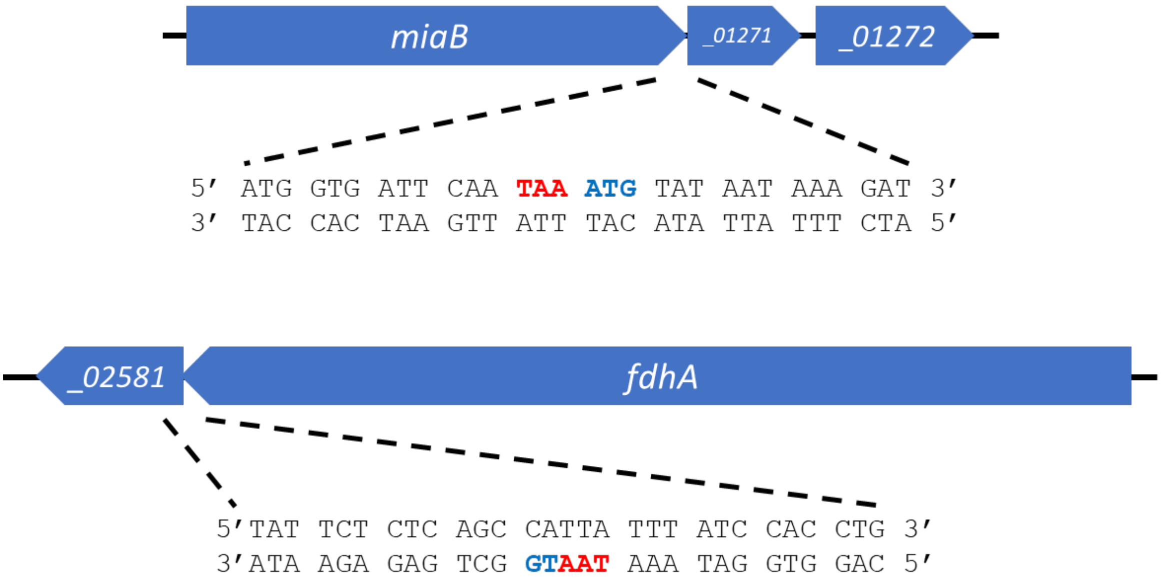

The *miaB* stop codon is directly adjacent to the *SAOUHSC_01271* start codon, while the *fdhA* stop codon overlaps the *SAOUHSC_02581* start codon. In both cases, the absence of a canonical Shine-Dalgarno sequence suggests translational coupling for the downstream gene.

### IsrR/target IntaRNA pairing predictions .Figure S3

| | | |
|---|---|---|
| **SAOUHSC_00304** | 1 19<br>\| \|<br>5'- GA GGUAU..GCC-3'<br>GGGUAUAUGAA GGGAAU<br>\|\|\|\|\|\|\|\|\|\|\| \|\|\|\|\|\|<br>**CCC**GUAUAUUU **CCC**UUA<br>3'-UGU..AAAUG U**C** UAAAC..GUU-5'<br>\| \|<br>96 78 | E= -15.54 kcal/mol<br><br>HE= -22.78 kcal/mol |
| **SAOUHSC_00410** | 22 38<br>\| \|<br>5'-AGA..ACAAA U AA AUAUA..AGU-3'<br>UAAA UG GU**GAGGG**U<br>\|\|\|\| \|\| \|\|\|\|\|\|\|\|<br>AUUU AC CAUU**CCC**A<br>3'-UGU..UUUGG C AG AAUGA..GUU-5'<br>\| \|<br>158 142 | E= - 7.14 kcal/mol<br><br>HE= -15.16 kcal/mol |
| **SAOUHSC_00545** | 79 86<br>\| \|<br>5'-AUA..AGAAA GCAAUA-3'<br>AUAAAACG<br>\|\|\|\|\|\|\|\|<br>UAUUUUGC<br>3'-UGU..GAAAA AACAA..GUU-5'<br>\| \|<br>130 123 | E= - 2.79 kcal/mol<br><br>HE= -6.18 kcal/mol |
| **SAOUHSC_01002** | 79 107<br>\| \|<br>5'-CCC..UCAAA AAAAUU UC U AUUA-3'<br>AAA**GUG**UC UAAG UUUGCU CU<br>\|\|\|\|\|\|\|\| \|\|\|\| \|\|\|\|\|\| \|\|<br>UUUCACAG AUU**C** AAAUGA GA<br>3'-UGU..UUGGA C **CC** C AAAUA..GUU-5'<br>\| \|<br>157 134 | E= - 9.72 kcal/mol<br><br>HE= -14.58 kcal/mol |
| **SAOUHSC_01437** | 90 96<br>\| \|<br>5'-CAG..UAUUC AUAAA..CCA-3'<br>G**GGGGG**A<br>\|\|\|\|\|\|\|<br>CCCCCCU<br>3'-UGU..AACAC ACAAG..GUU-5'<br>\| \|<br>38 32 | E= -10.55 kcal/mol<br><br>HE= -15.96 kcal/mol |
| **SAOUHSC_01960** | 77 87<br>\| \|<br>5'-GCC..AGCGU ACUAA..GCG-3'<br>GAAACGUU**GUG**<br>\|\|\|\|\|\|\|\|\|\|\|<br>CUUUGCAACA**C**<br>3'-UGU..UCUUG **CCCCC**..GUU-5'<br>\| \|<br>49 39 | E= -10.11 kcal/mol<br><br>HE= -15.28 kcal/mol |
| **SAOUHSC_02600** | 1 18<br>\| \|<br>5'- GUG**G** ACAAC..CAU-3'<br>AAGAGCGAAA **GUGG**<br>\|\|\|\|\|\|\|\|\|\| \|\|\|\|<br>UUCUUGCUUU CA**CC**<br>3'-UGU..GGUUG GCAA **CCCCC**..GUU-5'<br>\| \|<br>55 38 | E= -12.91 kcal/mol<br><br>HE= -19.14 kcal/mol |

IntaRNA (Mann et al., 2017) predictions for pairings between IsrR (lower sequence) and indicated mRNA (upper sequence). Blue bold sequences, SD; Black bold sequences, start codons; red bold sequences, CRRs; E, Energy; HE, Hybridization energy.

**Figure S4. 5'UTR reporter assay of IsrR putative targets.**

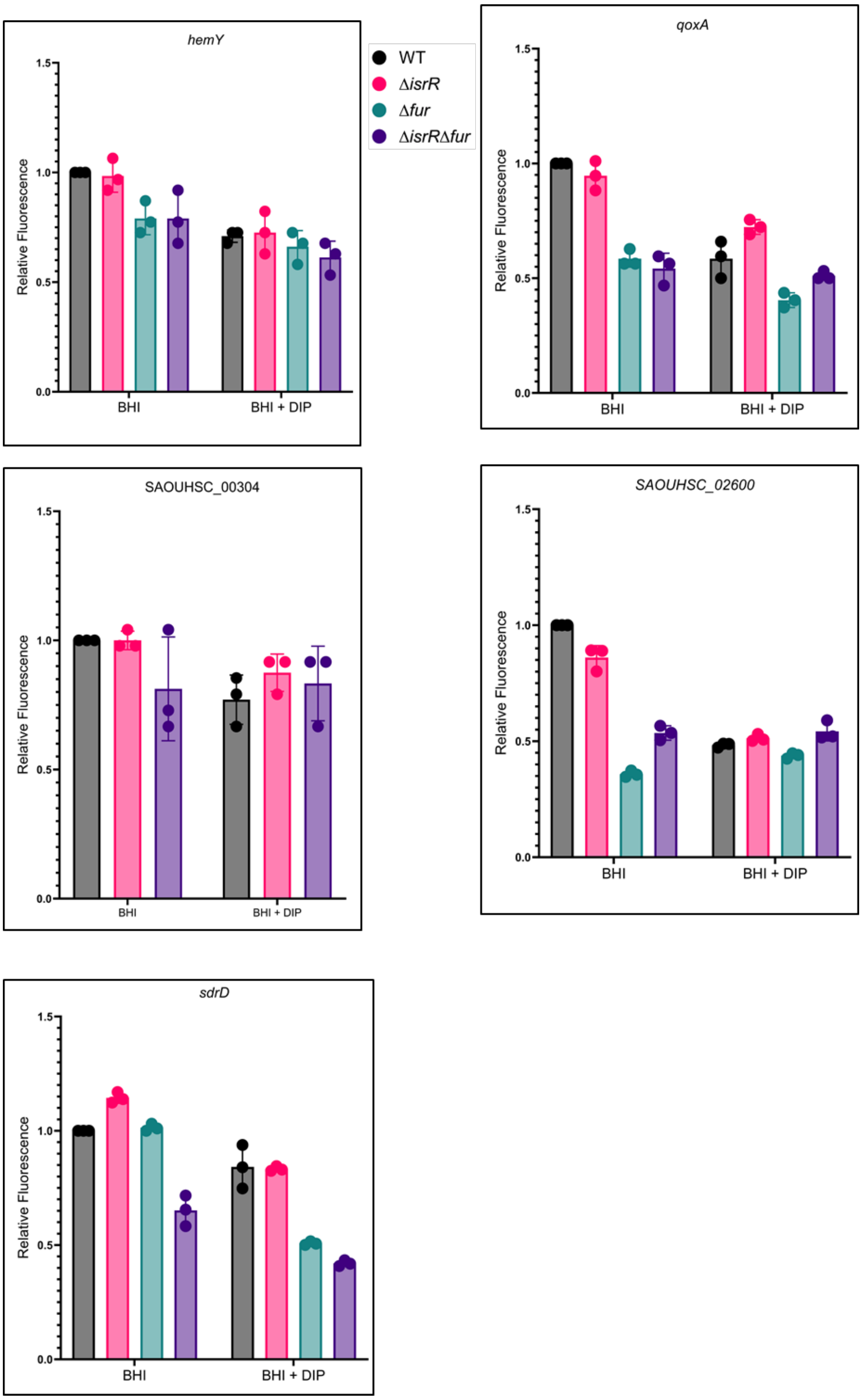

The 5'UTR–mAmetrine reporters of the indicated genes were integrated ectopically into the chromosome of *S. aureus* HG003 and its isogenic mutants (Δ*isrR*, Δ*fur*, and Δ*isrR* Δ*fur*). Cultures were grown in BHI medium with or without 2,2'-dipyridyl (DIP) supplementation. Relative fluorescence (normalized to parental strain in BHI) is shown as bar graphs (mean ± SD, *n* = 4). Note that IsrR is fully expressed in the absence of Fur or in the presence of DIP (Coronel-Tellez et al., 2022).

**Figure S5. The SAOUHSC_02924 (*gabT*) 5'UTR reporter is not regulated by RsaE under the tested conditions**

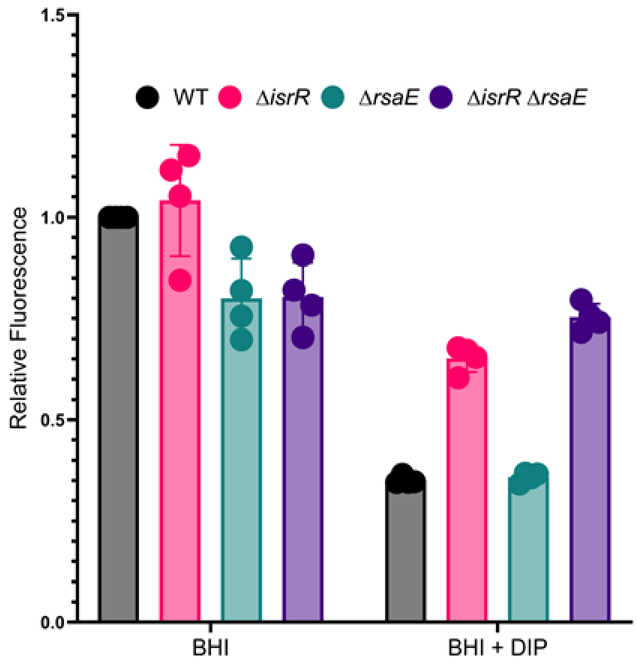

The *SAOUHSC_02924*–mAmetrine reporter was integrated ectopically into the chromosome of *S.aureus* HG003 and its isogenic derivatives (Δ*isrR*, Δ*rsaE*, and Δ*isrR* Δ*rsaR*) as indicated. Cultures were grown in BHI medium with or without 2,2'-dipyridyl (DIP) supplementation. Relative fluorescence (normalized to the parental strain in BHI) is shown as bar graphs (mean ± SD, *n* = 4).
